# Computational Analysis of Dynamical Fluctuations of Oncoprotein E7 (HPV) for the Hot Spot Residue Identification Using Elastic Network Model

**DOI:** 10.1101/401646

**Authors:** R. M. Malik, F. Nazir, S. Fazal, A. Bhatti, M. Ullah, S. I. Malik, A. Kanwal, S. E. Aziz, S. Azam

## Abstract

Virus proteins after invading human body alter host protein-protein interaction networks, resulting in the creation of new interactions, along with destroying or modifying other interactions or proteins. Topological features of new or modified networks compromise the host system causing increased production of viral particles. The molecular basis for this alteration of proteins interactivity is short linear peptide motifs similar in both virus and humans. These motifs are identified by modular domains, which are the subunits of a protein, in the human body, resulting in stabilization or moderation of these protein interactions Protein molecules can be modeled by elastic network models showing the fluctuations of residues when they are biologically active. We focused our computational study on the binding and competing interactions of the E7 protein of HPV with Rb protein. Our study was based on analysis of dynamic fluctuations of E7 in host cell and correlation analysis of specific residue found in motif of LxCxE, that is the key region in stabilizing interaction between E7 and Rb. Hot spot residue of E7 were also identified which could provide platform for drug prediction in future. Nevertheless, our study validates the role of linear binding motifs LxCxE of E7 of HPV in interacting with Rb as an important event in propagation of HPV in human cells and transformation of infection into cervical cancer.

## 1. Introduction

Our study was focused on analysis of interaction between viral oncoprotein E7 of Human Papilloma Virus (HPV) and human proteins, with use of computational methodology. Virus and cellular parasite proteins compete with host proteins for binding to mainitain the host protein-protein interaction (PPI) network, and alter these host interaction networks (Tournier & Quesnel-Hellmann, 2006; Sodhi et al., 2004; Dampier & Tozeren, 2007). Deep insight into the mechanism of interaction between host proteins and virus proteins provides a platform for drug discovery against virulance optimization of treatment for drugs already being in use (Brass et al., 2008). It is a challenge to study these interactions with the help of experiments due to involvement of about thirty thousand human proteins in proteome of a human (Roeth & Collins, 2006). Computational approaches have aided in experimental verification by reduction in the number of host proteins to be analyzed.

Searching for PPIs between viral and human proteins was the method on which the host-pathogen interaction prediction methods were focused previously. Advancement occured by the discovery of probability that human PPI network is operated by the interaction between two protein domains, and this probability was used to investigate the likelihood that human and viral proteins interact given their domain profiles (Dyer et al., 2007). In another method human and pathogen protein pairs were matched to proteins known to form complexes, and then these interaction candidates were filtered on the basis of expression data from pathogen and human (Davis et al., 2007). Due to presence of few domains on HPV proteins, and difficulty in finding their structures by comparative modeling, it is hard for scientists to transform these techniques to interactions between HPV and human proteins. For example, two different protein structures were required for comparative modeling to find structures for the N-terminal and C-terminal domains of HPV and host (Lv et al., 2007).

In present study, protein interactions interposed by short eukaryotic linear motifs (ELMs) on HPV proteins and human protein counter domains (CDs) known to interact with these ELMs were focused (Dinkel et al., 2016). A number of recent publications have discussed the potential functional roles of interactions mediated by ELMs and their CDs in viral infection (Tonikian et al., 2008; Shelton & Harris, 2008; Kadaveru et al., 2008). The HPV literature has shown the evidence that E7 for degradation of Rb tumour suppressor protein and related family members p107 and p130, it interacts with these protein through a conserved motif LXCXE at its amino terminal (Dyson et al., 1989; Munger et al., 1989). This experimental proof provides the motivation for systematic investigation of the association of motif/domain pairs with PPIs between virus and human proteins.

There is a well known characteristic of proteins is that they can exhibit various conformations in the neighborhood of their aboriginal conformation when folded, and these folded states are not rigid, but somewhat flexible (Frauenfelder et al., 1991). These can usually be considered as fluctuations near equilibrium positions, called equilibrium fluctuations. Two other more specific classes of conformational transitions accompany these fluctuations: 1) localized conformational isomeric movements sometimes occurring near native state coordinates, particularly for R groups with rotatable bonds; and 2) large-scale changes: functional equilibriumstates in some proteins that could be two or more depending on the function of protein (Damaschun et al., 1999; Frauenfelder & McMahon, 1998). Nevertheless, the most characteristic general fluctuations are usually small in magnitude, other than these two types, not exceeding few Ångstroms, and their frequency lies in the range of subnanosecond. By use of techniques developed for proteins based on normal mode analysis (NMA) and molecular dynamics (MD) with help of all atoms empirical potentials, we can elucidate details of molecular motions of proteins in the folded state (McCammon & Harvey, 1987; Brooks et al., 1988; Kitao & Go, 1999). Large size of the system make the techniques ineffective, involving atomic approaches with computational technology, even sometimes makes the result ambiguous. New advancements have developed coarse grained models and simplified force fields for description of vibrational dynamics of simple models, which has enabled the researchers to study proteins by applying them on overall dimensions of kinetics of complex systems or of the largest proteins, (Bahar et al., 1999; Bahar & Jernigan, 1999; Hinsen et al., 1999), consisting of several thousand or more residues. Conventional atomic models and potentials are not possible to use for the study of such systems. The dynamics of large numbers of proteins and their complexes within the scope of structural and functional study of genomes are systematically elucidated by pressing need of increasingly becoming important simple models and efficient computational methodologies (Amadei et al., 1993; van Aalten et al., 1997). It has been shown by a series of recent papers (Bahar et al., 1997, 1998a, 1999; Haliloglu et al., 1997; Bahar & Jernigan, 1998), that elastic networks model the fluctuation dynamics of proteins, in which residues are represented by nodes, and inter residual potentials stabilizing the folded conformation of proteins is shown as linkers. This model is also termed as the Gaussian network model (GNM) of proteins, in which Guassian distributed fluctuations are experienced by residues about their mean positions, also accompanied by harmonic potentials. Different types of amino acids are not distinguishable, due to which all residues are to be considered to adopt a generic force constant for the interaction potential between all residue pairs close enough. New neutron sacattering experiments are in laboraotories to generate experimental values of these generic force constants, and that will provide a more precise method to find the values of these constants (Zaccai, 2000). The magnitudes of fluctuations have been verified to be provided by GNM. In the GNM there is no consideration of directions or 3-D characterisitics of movements of residues, all fluctuations are unexpressively assumed to be isotropic. Each residue is represented as N and protein molecule as a cluster of N sites, forming an ensemble of N having 1 independent mode. No involvement of 3D description which would show 3 N 6 modes instead of 1. It is necessary to keep in view the directions of collective movements of residues as they play a major role in biological functioning and mechanisms of interactions of proteins, so regarding the proteins, these fluctuations are anisotropic instead of being isotropic (Kuriyan et al., 1986; Ichiye & Karplus, 1987). It is not indeed possible to acquire an understanding of the mechanism of motion unless the fluctuation vectors, in addition to their magnitudes, are elucidated. This issue has been addressed by an advanced extension of the GNM, called the anisotropic network model (ANM). The development of the ANM has occured as an extension of a recent comparison of the results from MD simulations with the GNM for treating the anisotropy of fluctuations (Doruker et al., 2000). The elastic network model is a coarse-grained approach that is used for investigating dynamics of protein machinary (Tirion, 1996; Bahar et al., 1997). In these models single identical beads typically represent all amino acid residues of the protein, while empiric harmonic potentials depending only on the distance between any two beads mediate the interactions between them. A protein is studied as an elastic object in these models, having a network of beads connected through distortable elastic springs. These models have been validated being powerful in describing ATP-induced slow motions in molecular machines and advanced the understanding of important aspects of their collective dynamics, despite the coarse-grained nature of elastic network descriptions and the gross simplification made (Hinsen, 1998; Bahar et al., 1998; Tama & Sanejouand, 2001; Zheng & Doniach, 2003; Yang et al., 2007; Flechsig et al., 2011). Our target was to analyze host pathogen proteins interactions at residual level based on ELM and CD associations, alongwith the elastic network modelling of viral proteins so that we have an insight into the flexible regions of viral proteins and their role in establishing these interactions.

## Methods

The protein sequences of E7 protein were downloaded from UniProt (Pundir et al., 2017), having ID No.P03129. E7 was annotated with ELMs using the ELM resource (Dinkel et al., 2016), using default settings except selecting the human for the species field. We got the list of ELMs found in the E7 matching the sequences of motifs in humans.

By using version 2016 of NMA tool ANM 2.1 (Eyal, 2015) the elastic networks of the E7 protein were modeled. A 3D structure of E7 was also constructed by using the web portal RaptorX (Ma et al., 2013), by giving FASTA sequence of E7 downloaded from UniProt. This 3D structure was used as input to ANM2.1.

Elastic network models and graphs showing the fluctuations of amino acids in 20 different modes for E7 were used for analysis of interactions of E7 with Rb protein.

## 2. Results and Discussion

E7 is a protein composed of 98 amino acids (Dyson et al., 1989; Munger et al., 1989). Among its amino acids sequence there are 18 regions which possess sequences similar to the sequences of different linear motifs found in human proteins. As motifs are very important and play a major role in developing an interaction between proteins. For stabilizing these inetractions the motifs develop a link with specific sequences in other proteins called domains. The motifs were selected on the basis of probability of p<0.1. These motifs are enlisted in table 1, alongwith the domains with which they interact.

**Table 1:** Motifs in E7 with their Counter Domains.

| S.No. | ELMs | Counter Domains (CDs) |
| --- | --- | --- |
| 1. | CLV_C14_Caspase3-7 | Peptidase_C14 |
| 2. | DOC_MAPK_MEF2A_6 | Pkinase |
| 3. | MOD_CK2_1 |  |
| 4. | MOD_NEK2_1 |  |
| 5. | MOD_NEK2_2 |  |
| 6. | MOD_ProDKin_1 |  |
| 7. | MOD_PLK |  |
| 8. | DOC_USP7_MATH_1 | MATH |
| 9. | LIG_TRAF2_1 |  |
| 10. | DOC_WW_Pin1_4 | WW |
| 11. | LIG_FHA_1 | FHA |
| 12. | LIG_FHA_2 |  |
| 13. | LIG_NRBOX | Hormone_recep |
| 14. | LIG_Rb_LxCxE_1 | RB_B |
| 15. | LIG_SH2_STAT5 | SH2 |
| 16. | MOD_N-GLC_1 | STT3 |
| 17. | MOD_PIKK_1 | PI3_PI4_kinase |
| 18. | TRG_ENDOCYTIC_2 | Adap_comp_sub |

### 2.1 ELASTIC NETWORK MODELS OF E7

Elastic network models were generated in 20 different modes of protein E7. The elastic network models are shown in figure 1. The protein is shown to exhibit possible movement in respective mode. Red coloured regions of protein chain represent the residue which show the maximum degree of fluctuation in each mode.

**Figure 1.**
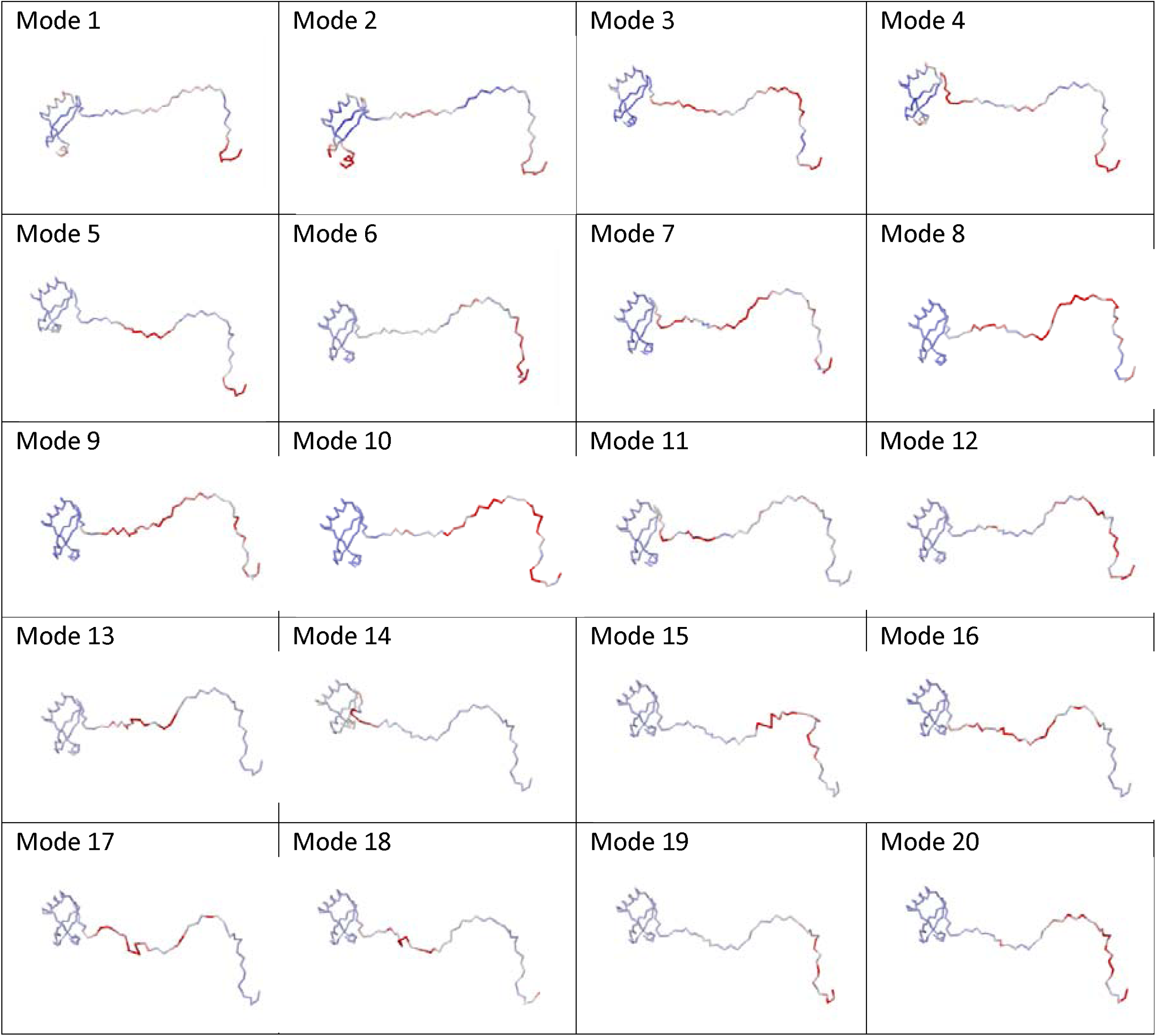
The elastic network models of E7 are shown in 20 different modes. The red colour of chain represents the maximum degree of flexibility of chain and fluctuation of amino acid residues lying in that highlighted region.

Mode 1: In mode 1 of E7 the amino acids which appear to show maximum flexibilty and fluctuation are situated between position 1 and 10. This region contains the sequences similar to those of motifs MOD_ProDKin_1, DOC_WW_Pin_1_4, MOD_CK2_1 and LIG_FHA_2.

Mode 2: The regions showing maximum degree of fluctuation and flexibility in mode 2 lie between the positions 55-70 and 91-98. Motifs to be found in these regions are DOC_MAPK_MEF2A_6, LIG_FHA_1, MOD_NEK2_2, MOD_PLK, MOD_PIKK_1 AND DOC_USP7_MATH_1.

Mode 3: The fluctuating regions in mode 3 possess sequence/s for the motifs of CLV_C14_Caspase 3-7, DOC_WW_Pin1-4, LIG_FHA_1, LIG_FHA_2, LIG_Rb_LxCxE_1, LIG_SH2_STAT5, LIG_TRAF2-1. MOD_CK2_1, MOD_N_GLC_1, MOD_ProDkin_1 and TRG_ENDOCYTIC_2.

Mode 4: In mode 4, E7 shows 2 regions having higher degree of fluctuation, one lies in the region same as in mode 1 at positions 1-10 and other at positions 37-55 which possesses the sequences for the motifs of CLV_C14_Caspase 3-7 and TRG_ENDOCYTIC_2.

Mode 5: Mode 5 exhibits flexibility on positions 1-10 and 25-40. These regions show similarity to the sequence/s of motifs DOC_WW_Pin1_4, LIG_FHA_1, LIG_FHA_2, LIG_Rb_LxCxE_1, LIG_SH2_STAT5, LIG_TRAF2_1, MOD_CK2_1, MOD_N-GLC_1, MOD_ProDKin_1, and TRG_ENDOCYTIC_2.

Mode 6: In mode 6 of E7 the amino acids which appear to show maximum flexibilty and fluctuation are situated between position 1 and 10, like that of mode 1. This region contains the sequences similar to those of motifs MOD_ProDKin_1, DOC_WW_Pin_1_4, MOD_CK2_1 and LIG_FHA_2.

Mode 7: The amino acids showing highest degree of fluctuation in mode 7 at position 19-55. The motifs lying in these regions include CLV_C14_Caspase3-7, LIG_FHA_1, LIG_Rb_LxCxE_1, LIG_SH2_STAT5, LIG_TRAF2_1, MOD_CK2_1, MOD_N-GLC_1, and TRG_ENDOCYTIC_2. Mode 8: The mode 8 shows fluctuation in region of 19-28 positions of amino acids. These sequences match with the sequences of LIG_FHA_1, LIG_Rb_LxCxE_1, and LIG_SH2_STAT5.

Mode 9: The most flexible region of mode 9 lies in the region of amino acids situating at between positions 35 and 42. These sequences possess the similarity with motif CLV_C14_Caspase3-7.

Mode 10:The most flexible region exihibited by the amino acids in mode 10 is lying from positions 4-10. It possesses the sequences of motifs LIG_FHA_1, LIG_FHA_2 and MOD_CK2-1.

Mode 11: The amino acids showing maximum degree of movement in mode 11 are from position 32-40, with the possibility of presence of motifs CLV_C14_Caspase3-7 and LIG_TRAF2_1.

Mode 12: The mode 12 shows the amino acids from positions 14-23 to be most flexible having the motif sequences of LIG_FHA_1 and LIG_Rb_LxCxE_1.

Mode 13: LIG_TRAF2_1 shows its presence in the mode 13 due to flexibility of the amino acids at positions 31-35.

Mode 14: No motif has been found matching the sequence of amino acids lying in the most flexible region of the mode 14 at positions 40-45.

Mode 15: Amino acids showing the maximum degree of fluctuation in mode 15 are from position 23-28. Motifs having such sequences are LIG_SH2_STAT5 and TRG_ENDOCYTIC_2.

Mode 16: In mode 16 two regions are showing flexibility, one from position 22-26 and other 28-35. Motifs lying in these rgions are LIG_Rb_LxCxE_1, LIG_SH2_STAT5, LIG_TRAF2_1, MOD_CK2-1, MOD_N-GLC_1, and TRG_ENDOCYTIC_2.

Mode 17: LIG_TRAF2_1 is the only motif showing its presence in the mode 17 due to maximum fluctuation exhibited by the amino acids of the sequence identical to it, at positions 32-37.

Mode 18: There are 2 flexible regions in the mode 18 one at position 30-32 and other at 33-37. LIG_TRAF2_1 and MOD_CK2-1 show similarity to the amino acids sequences present in these regions.

Mode 19: The mode 19 is occupied by three motifs LIG_FHA_2, MOD_CK2-1 and MOD_ProDKin_1 being similar in amino acid sequence at position 5-10. Amino acid 1 shows the highest value of fluctuation in this mode.

Mode 20: In mode 20 amino acids at positions22-26 show the maximum flexibility and their sequence is similar to those of motifs LIG_SH2_STAT5 and TRG_ENDOCYTIC_2.

### 2.2 Observed Conformational changes in E7 for interaction with Rb tumor Suppressor protein

Modes 3, 5, 7, 8, 12 & 16 are the the modes showing fluctutations in residues forming the LxCxE motif that is involved in interaction with Rb protein, as shown in figure 2. The retinoblastoma (Rb) protein family has been found to be interacted by multiple viral proteins through the LxCxE motif. The B domain of Rb protein possesses a highly conserved and shallow groove, that is involved in its interaction with E7, mediated through the LxCxE motif. The central Cysteine is highly conserved in all instances, however, the Leucine and Glutamic Acid positions tolerate substitution of physicochemically similar residues allowing a less stringent definition of [LI]xCx[DE] (Liu & Marmorstein, 2007). All these modes show maximum degree of fluctuation in the region of E7 contraining sequence of motif LxCxE, as depicted by the sequence in flexible region of residues 21-37, i.e. DLYCYEQLNDSSEEEDE. The staggered arrangement, evenly spaced and one residue apart, of the conserved residues cover one side of an extended, beta-strand-like conformation and bind the groove orthogonally, not by beta augmentation like many similar staggered motifs. The Leucine and Cysteine positions bind a hydrophobic region of the groove with tight complementarity. The Glutamic Acid forms hydrogen bonds with two backbone amide groups of an alpha helix forming one side of the binding groove. The interaction is further stabilized by additional hydrogen bonds to the peptide backbone adding rigidity. Phosphorylation of Rb at Thr821 and Thr826 inhibits LxCxE binding. Fig 21 to 22 show the degree of flexibility of this region of E7 in different modes.

**Figure 2.**
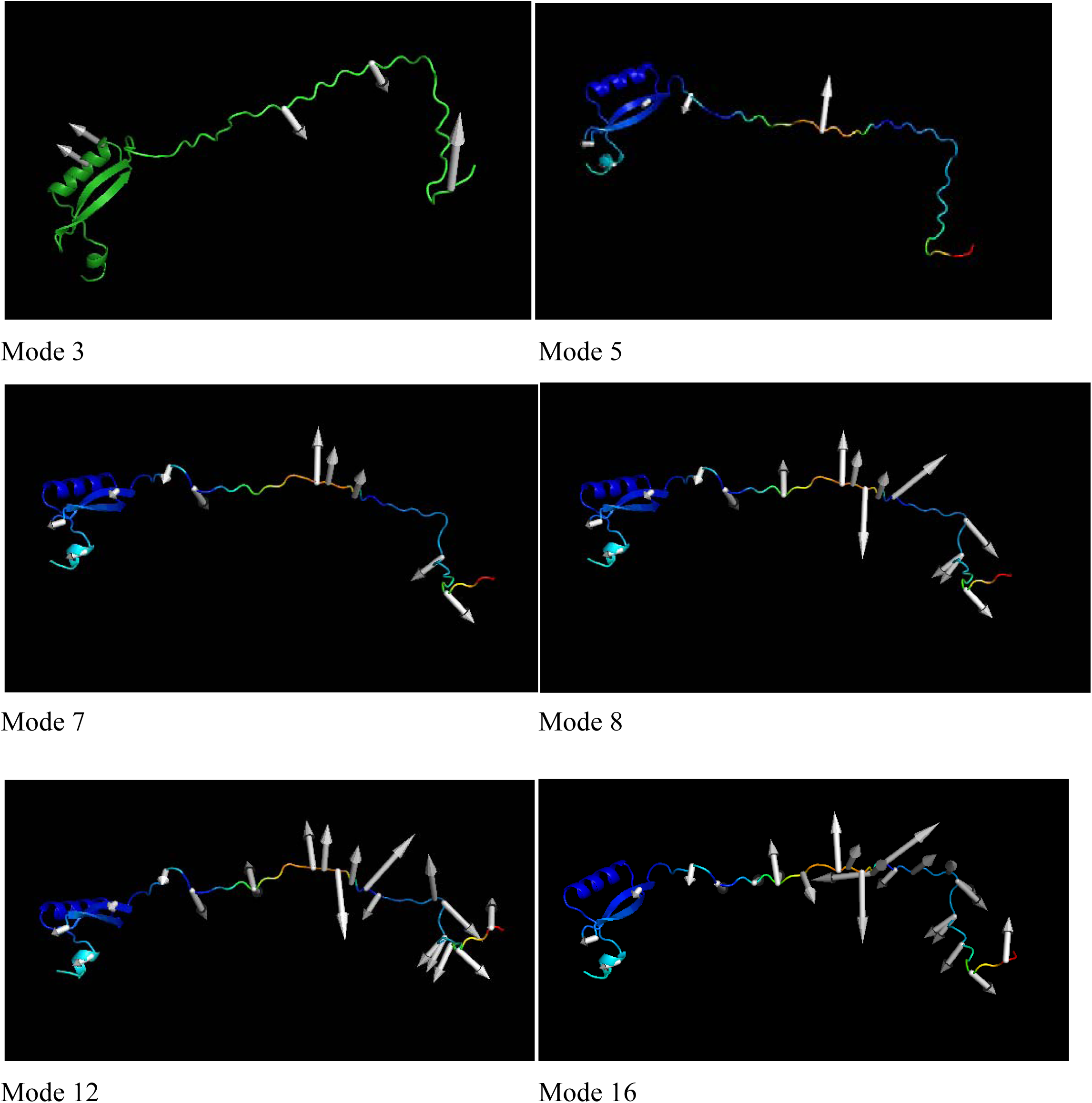
the selected modes (3, 5, 7, 8, 12 & 16) of E7 are shown as elastic network models. The arrows are showing the degree of fluctutaion of specific regions. The area of interest for us is the regions which contain sequence for the LxCxE motif

### 2.3 Distance weights for force constants

The plots of B factors of 20 modes of E7 are shown in Fig.3. Graphs of Mode 3, 5, 7, 8, 12 & 16 show highest B factors in regions of motif LxCxE (Hinsen 1998). Which confirms their involvement in stabilizing the interaction with Rb protein.

**Figure 3.**
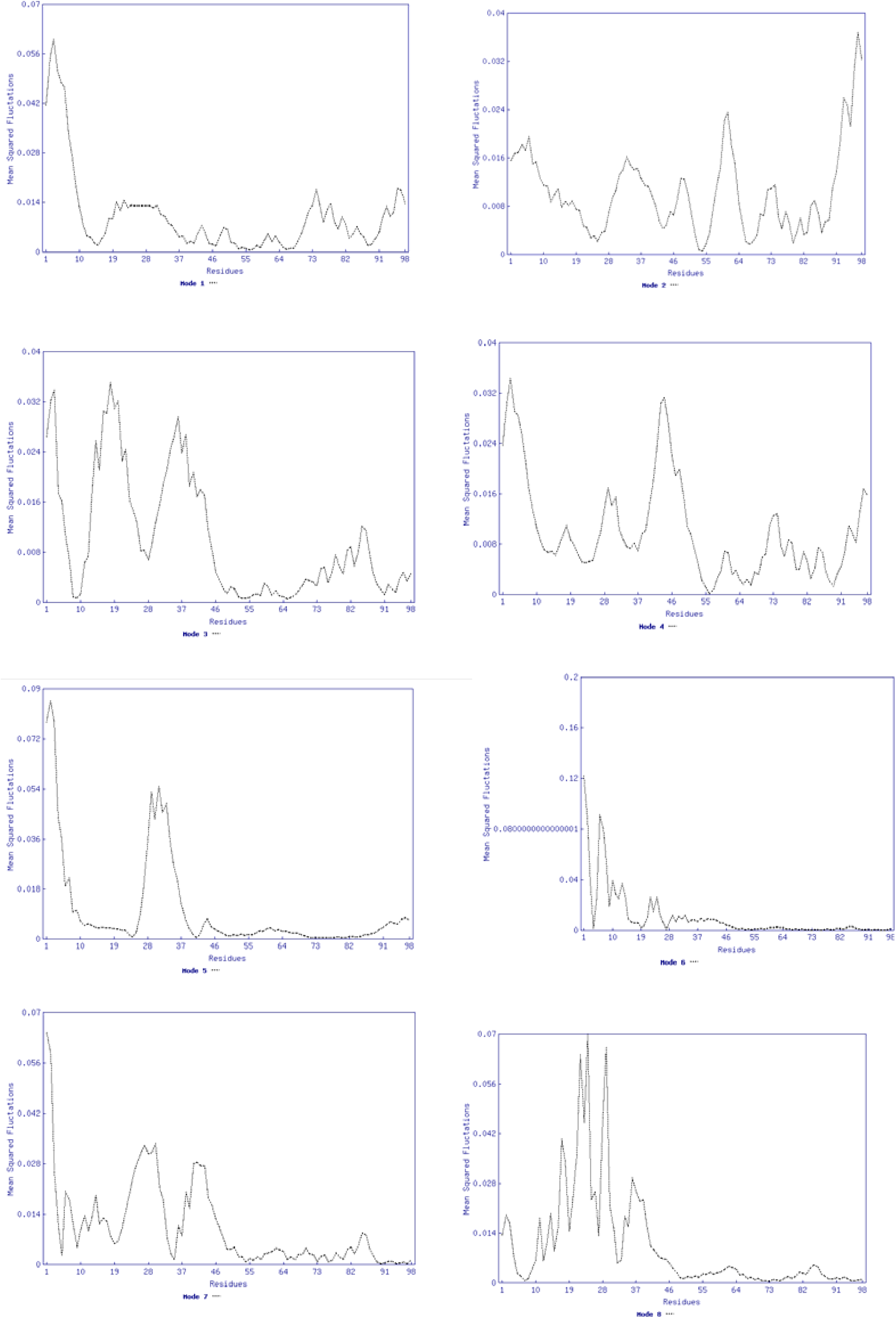
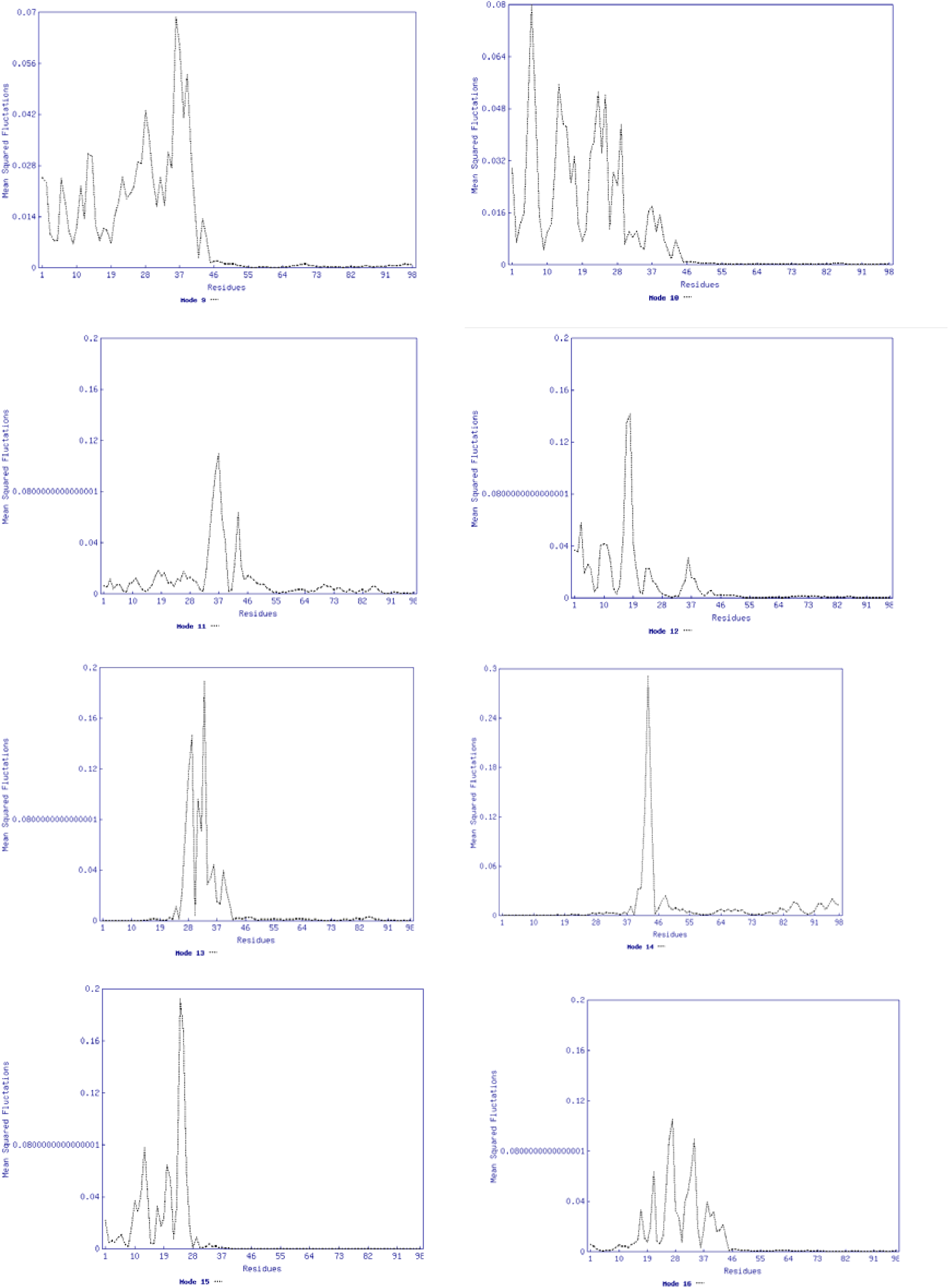
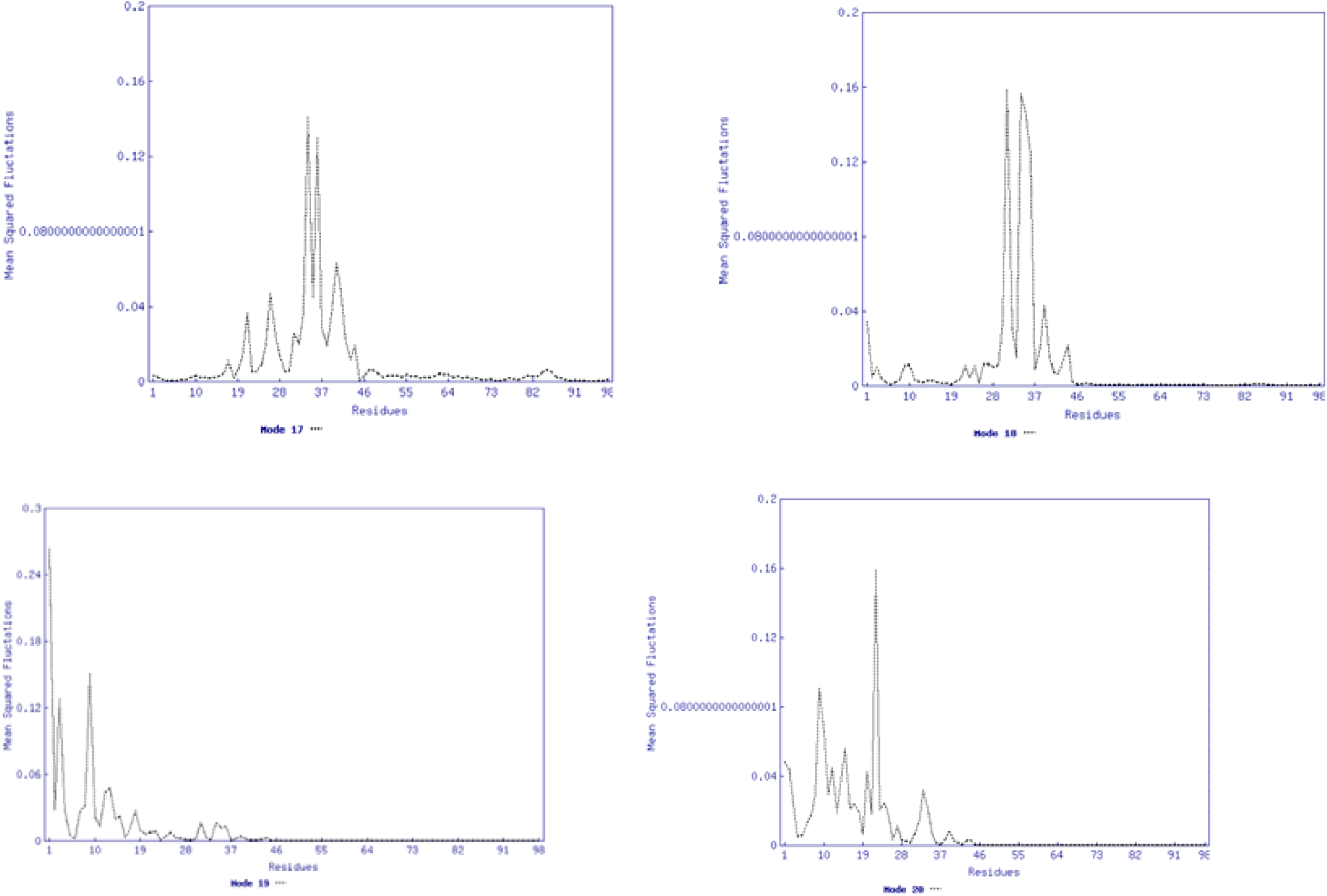
Graphs showing the B factors of amino acid residues of E7 in all 20 different modes.

### 2.4 Correlation Analysis of E7

As it is very well explored elsewhere (Bahar et al., 1997, Atilgan et al., 2001, Hinsen et al., 1999, Chen & Bahar, 2004, Brooke et al., 1988 and Sen et al 2006), the dynamics of the protein structures can be explained well by NMA of an ENM. The important motions of proteins are characterized by the lowest frequency normal modes which are universally the most dominant. For further investigation, each pair of residues was considered for the correlations of the motions. Correlation analysis is able to detect and capture the motif LxCxE motions in stabilizing the interaction with Rb protein. We used The standard ANM approach, which analyzes co-ordinately moving blocks of amino acid residues, and gives information directly about their correlations, for each mode (Atilgan et al., 2001). In fig 4. the correlation plots for modes (3, 5,7, 8, 12 & 16) are depicted. In these modes we again experience very high correlations of the LxCxE motif region, giving evidence of their fluctuations and invovlement in maintaining interactions with Rb protein.

**Figure 4.**
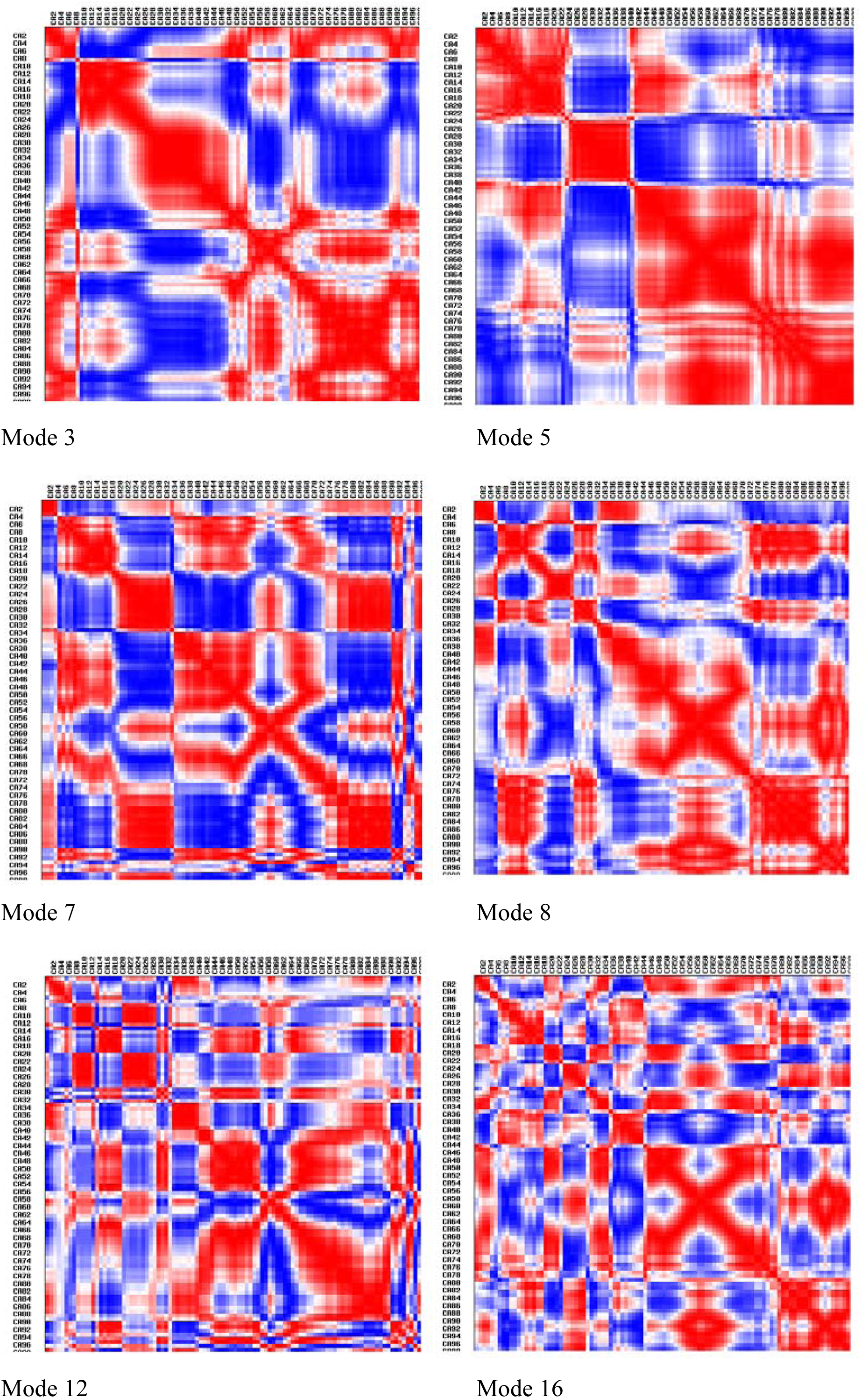
Correlation for E7 protein. The color scheme of matrix is such that large inter-residue fluctuations appear as blue and small fluctuations as red. Specific Blue regions show correlation of residues of LxCxE motif with different amino acid residues.

**Figure 5:**
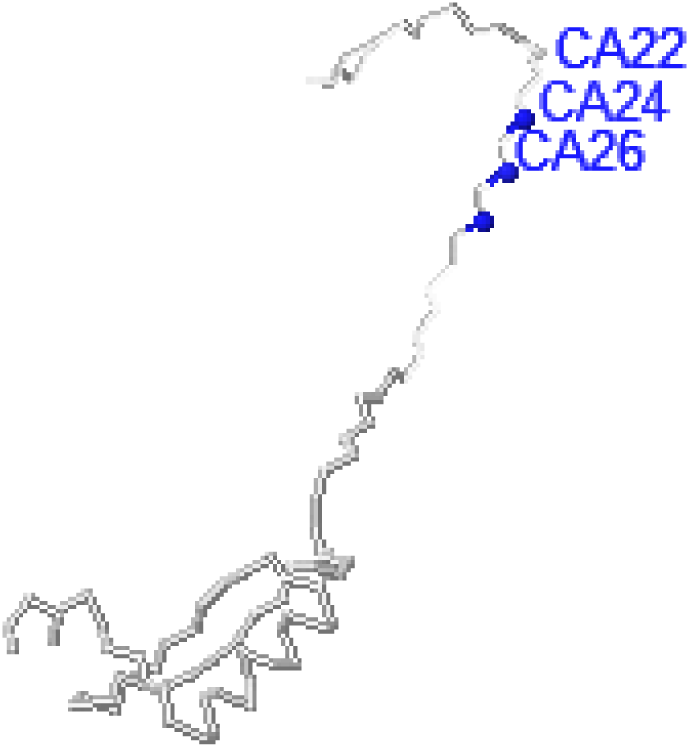
The hot spot residues of E7, which play a vital role in interaction of E7 with Rb protein.

On the basis of above findings of dynamics fluctuations of E7, for the subset of Rb binding residues, we find that its hot-spot residues, whose perturbations lead to a significant changes in the fluctuations of Rb-binding residues, span an extensive network that not only interacts with Rb protein, but also stabilizes the interaction to create the cascade required in response. L 22, C24 and E26 of LxCxE motif are highly conserved residues of E7, which are found to be dynamically involve in interaction with Thr821 and Thr826 of Rb protein. These key residues are proposed to mediate dynamical couplings and mutual modulations between the E7 and Rb protein.

## Conclusion

In this study we have investigated the dynamical mechanisms in E7 based on a simple elastic model. On the basis of the presence of a specific motif region in E7, that is involved in its interaction with Rb protein, in order to cause carcinogenicity by HPV, we have found the six different modes to dominate the observed changes, thus validating the application of ENM in E7. Using our ENM-based correlation analysis we have identified dynamically important hot spot residues, which are consistent with published evidences of involvement of specific amino acid residues in interaction of E7 with Rb protein.These hot spots could be used as drug targets for prevention of development of HPV infection into cancer. This provides suggestions for future experiments. Our correlation analysis provides a dynamic model for the binding regions of E7 with Rb protein, which provides evidence for the mechansim reported about this interaction (Dyson et al., 1989; Munger et al., 1989).

Computer representation of receptor flexibility is a major consideration in the de novo design of small molecules inhibitors (Alberts et al., 2005). In particular, the ability to represent backbone changes taking place during allosteric motions has so far been unavailable. The demonstration of the potential of the ENM methods presented in this paper provides a promising approach, which will allow representations of backbone conformational changes whose consideration is important for drug-design purposes.

